# Estimating the implicit component of visuomotor rotation learning by constraining movement preparation time

**DOI:** 10.1101/082420

**Authors:** Li-Ann Leow, Reece Gunn, Welber Marinovic, Timothy J Carroll

## Abstract

When sensory feedback is perturbed, accurate movement is restored by a combination of implicit processes and deliberate re-aiming to strategically compensate for errors. Here, we directly compare two methods used previously to dissociate implicit from explicit learning on a trial-by-trial basis: 1) asking participants to report the direction that they aim their movements, and contrasting this with the directions of the target and the movement that they actually produce, 2) manipulating movement preparation time. By instructing participants to re-aim without a sensory perturbation, we show that re-aiming is possible even with the shortest possible preparation times, particularly when targets are narrowly distributed. Nonetheless, re-aiming is effortful and comes at the cost of increased variability, so we tested whether constraining preparation time is sufficient to suppress strategic re-aiming during adaptation to visuomotor rotation with a broad target distribution. The rate and extent of error reduction under preparation time constraints were similar to estimates of implicit learning obtained from self-report without time pressure, suggesting that participants chose not to apply a re-aiming strategy to correct visual errors under time pressure. Surprisingly, participants who reported aiming directions showed less implicit learning according to an alternative measure, obtained during trials performed without visual feedback. This suggests that the process of reporting can affect the extent or persistence of implicit learning. The data extend existing evidence that restricting preparation time can suppress explicit re-aiming, and provide an estimate of implicit visuomotor rotation learning that does not require participants to report their aiming directions.

**New and Noteworthy:** During sensorimotor adaptation, implicit, error-driven learning can be isolated from explicit strategy-driven re-aiming by subtracting self-reported aiming directions from movement directions, or by restricting movement preparation time. Here, we compared the two methods. Restricting preparation times did not eliminate re-aiming, but was sufficient to suppress reaiming during adaptation with widely-distributed targets. The self-report method produced a discrepancy in implicit learning estimated by subtracting aiming directions, and implicit learning measured in no-feedback trials.

## Introduction

When we move, perturbations to our body or the environment can elicit discrepancies between predicted and actual outcomes. We readily adapt our movements to compensate when such discrepancies are systematic, and this process is commonly termed sensorimotor adaptation. Sensorimotor adaptation was traditionally thought to occur largely via implicit mechanisms involving updating of an internal model (Wolpert et al. 1995) in order to compensate for sensory prediction errors (i.e. mismatches between predicted and observed behaviour). It has long been recognized, however, that explicit processes can influence the behavioural response to sensorimotor perturbation (e.g., Keisler and Shadmehr 2010; Mazzoni and Krakauer 2006; Redding and Wallace 1996; Uhlarik 1973). For example, if a rotation of visual feedback results in a participant noticing systematic reaching errors to one side of a target, she might deliberately aim to the opposite side of the target to compensate. One way to disentangle such strategic re-aiming from implicit learning is to require participants to report their aiming directions throughout adaptation, and then to infer implicit adaptation by subtracting verbally reported aiming directions from actual movement directions (Bond and Taylor 2015; Brudner et al. 2016; McDougle et al. 2015; Taylor et al. 2014). This method also provides a measure of explicit re-aiming, which is estimated as the difference between the reported aiming direction and the target direction. Studies using this approach suggest that explicit re-aiming dominates the rapid initial error reduction typically seen in most sensorimotor adaptation studies, but then contributes progressively less to behaviour as an implicit *remapping* between motor commands and expected sensory outcomes develops with extended exposure to perturbation.

The capacity to decompose sensorimotor adaptation into implicit and explicit components represents an important advance in the understanding of how the brain responds to systematic discrepancies between desired and actual motor behaviour (Taylor et al. 2014). In particular, the demonstration that explicit re-aiming dominates the initial error reduction phase of sensorimotor adaptation presents challenges for those interested in assessing the rate of implicit remapping. A method of disentangling explicit and implicit processes that relies upon subject reports of aiming directions may have limitations, however. Firstly, the approach requires faithful reports of intended aiming directions from study participants, which may be imprecise, difficult to obtain in some contexts, and time-consuming. Secondly, the instruction to report aiming directions results in faster error reduction than occurs in the absence of such instructions (Taylor et al. 2014), presumably because the reporting requirement alerts participants to the benefits of re-aiming to achieve task success. This raises the question of whether the reporting procedure might also impact implicit processes, because the reduced task errors that accompany explicit strategy use might affect the rate or extent of implicit adaptation via reward or reinforcement-related processes (Reichenthal et al. 2016).

An alternative approach to probe implicit processes in sensorimotor adaptation is to suppress the expression of explicit processes. This can be done either by employing dual-task paradigms to limit attentional resources that can be devoted to explicit re-aiming (Galea 2010; Keisler and Shadmehr 2010; Malone and Bastian; Taylor and Thoroughman 2007; Taylor and Thoroughman 2008), or by restricting the amount of time available to prepare a movement (Fernandez-Ruiz et al. 2011; Haith et al. 2015). Restricting preparation time appears to be a particularly promising approach, as there is a relationship between preparation time and movement accuracy even without a sensorimotor perturbation (Georgopoulos and Massey 1987b; Marinovic et al. 2017). Furthermore, there is a time cost of explicitly preparing movements toward locations that are offset from the physical location of a target (Georgopoulos and Massey 1987b). In one such approach, Haith et al. (2015) carefully controlled movement preparation time to dissociate learning resulting from explicit and implicit processes during adaptation to a visuomotor rotation. They showed significantly slower error reduction when they restricted movement preparation time by suddenly shifting target position in 20% of trials approximately 300ms before the imperative to move (Haith et al. 2015). The data suggest that explicit re-aiming was supressed by the preparation time constraint. The approach also has the benefit that it provides a within-subject contrast between presumed implicit remapping (from errors on the short preparation trials) and combined implicit and explicit adaptation (from errors on the long preparation trials). However, some aspects of this approach merit further consideration. First, it is unclear whether 300 ms is sufficiently brief to prevent entirely strategy use during adaptation. Second, the switch in target location might introduce an additional processing demand, and may not be desirable in some experimental designs. More generally, it is unknown whether assays of implicit sensorimotor adaptation obtained via preparation time manipulation differ from those obtained via reporting procedures. Here, we compared implicit learning assayed by restricting movement preparation time to implicit learning assayed via reporting procedures.

The first aim of the study was to determine the extent to which the capacity to explicitly re-aim is suppressed by reducing the amount of time available to prepare movement. We asked people to explicitly re-aim 30° clockwise or counter-clockwise to targets, under increasing time pressure, but in the absence of a perturbation. We expected that there would be a minimum time for movement preparation below which people would be unable to aim accurately to one side of a target. However, we also wondered whether advance knowledge of the approximate location of potential targets would influence the capacity to re-aim. To this end, voluntary re-aiming was performed either to a narrow (0-35° range) (Experiment 1A) or uniform 360°distribution of target directions (Experiment 1B). We predicted that people would be able to re-aim with shorter preparation times when targets were distributed narrowly. We found that participants could re-aim by 30° even at the shortest preparation times tested with a narrow target distribution, but at the expense of increased movement variability. For a broad 360° target distribution, participants could at least partially re-aim whenever movement time was sufficient to produce directionally tuned movements (i.e., as opposed to randomly directed movements), but at more dramatic cost to movement variability. Thus, the motor system is capable of systematic re-aiming to one side of a target irrespective of time constraints. However, we noted that participants found re-aiming at short preparation times extremely effortful. Given this, the purpose of Experiment 2 was to determine whether people would choose to re-aim under time pressure in order to improve performance on a visuomotor rotation task.

In Experiment 2, we compared adaptation to a 30° visuomotor rotation with a 360° target distribution under three alternative conditions. Separate groups of participants were either allowed: (1) a short time to prepare movement, (2) a longer time to prepare movement, but also asked to report their aiming direction, or (3) a longer time to prepare movement, without reporting aiming direction. If people chose not to re-aim reaches to counter the visuomotor rotation when preparation time was constrained, then we expected the rate of error reduction in this condition to resemble the rate of implicit adaptation estimated from the self-report procedure. We were also interested in the effects of the three different conditions on an alternative measure of implicit adaptation obtained from reaches made in the absence of visual feedback. We found that the rate and extent of error compensation with short preparation time closely matched implicit error compensation, as estimated from subtracting movement directions from self-reported aiming directions. This suggests that restriction of preparation time can suppress explicit re-aiming, and provide an estimate of implicit learning that does not require participants to report their aiming directions. Surprisingly, in the post-perturbation no-feedback trials, less implicit learning was shown in participants who reported aiming directions than participants who did not report aiming directions. This raises the possibility that the reporting procedure itself increased engagement of explicit learning, which inadvertently reduced engagement of implicit learning.

## Method

### Participants

A total of 74 participants completed this study (Experiment 1A: n=14, mean age = 19.93, range = 17-42 years, 12 females, 2 left-handed; Experiment 1B: n=14, mean age = 19.07, SD = 3.53, range = 17-31 years, 11 females, 2 left-handed; Experiment 2: n=36, 30 females, 2 left handed, mean age =19.85, SD = 1.82). In Experiment 2, 36 people were initially assigned either to a short preparation time condition or a long preparation time condition in which they had to report aiming direction. Subsequently, in order to test whether differences in post-perturbation estimates of implicit learning were due to the preparation time conditions or the reporting procedure, a further 10 people were recruited to a long preparation time condition without reporting (mean age 21, SD=4.7, range=18 to 34 years, all right-handed). For all experiments, the participants were randomly assigned either to clockwise or counter-clockwise visuomotor rotation conditions in equal proportions. All participants were naïve to visuomotor rotation and force-field adaptation tasks.

### Apparatus and General Trial Structure

Participants completed the task using the vBOT planar robotic manipulandum, which has a low-mass, two-link carbon fibre arm and measures position with optical encoders sampled at 1,000 Hz (Howard et al. 2009). Participants were seated on a height-adjustable chair at their ideal height for viewing the screen for the duration of the experiment. Visual feedback was presented on a horizontal plane on a 27” LCD computer monitor (ASUS, VG278H, set at 60Hz refresh rate) mounted above the vBOT and projected to the subject via a mirror in a darkened room, preventing direct vision of their hand. The mirror allowed the visual feedback of the target (a 0.5 cm radius circle), the starting location (a 0.5 cm radius circle), and hand cursor (0.25 cm radius) to be presented in the plane of movement, with a black background. The start circle was aligned 10cm to the right of the participant’s mid-sagittal plane at approximately mid-sternum level.

### General Trial Structure

Participants made centre-out reaching movements by moving the robot arm from the start circle to the target. Targets appeared in random order at one of eight locations 9cm away from the start circle—target locations were clustered either in a small range (Experiment 1A: 17.5°,12.5°,7.5°,2.5°,−2.5°,−7.5°,−12.5°,−17.5° from straight ahead), or distributed uniformly throughout 360° (Experiment 1B & Experiment 2: 0°, 45°, 90°, 135°, 180°, 225°, 270° and 315°). At the start of each trial, the central start circle was displayed. If participants failed to move the hand to within 1cm of the start circle after 1 second, the robotic manipulandum passively moved the participant’s hand to the start circle (using a simulated 2 dimensional spring with the spring constant magnitude increasing linearly over time). A trial was initiated when the cursor remained within the home location at a speed below 0.1 cm/s for 200 ms. We used a timed-response paradigm (Ghez et al. 1989; Haith et al. 2015; Marinovic et al. 2014; Marinovic et al. 2008; Schouten and Bekker 1967) to manipulate movement preparation time. Across all conditions, a sequence of three tones spaced 500 ms apart was presented at a clearly audible volume via external speakers. Participants were instructed to time the onset of their movements with the onset of the third tone (see Figure 1). They were instructed not to stop on the target, but to slice through it. Movement initiation was defined online as when hand speed exceeded 2cm/s. Targets appeared at 1000ms, 250ms, 200ms, 150ms, or 100ms, minus a display latency (27.6 ± 1.8 ms), prior to the third tone. Thus target direction information became available 972.4, 222.4, 172.4, 122.4, or 72.4 ms before the desired initiation time. When movements were initiated 50 ms later than the third tone, the trial was aborted: the screen was blanked and a “Too Late” on-screen error signal appeared. Similarly, when movements were initiated more than 100 ms before the desired initiation time, the trial was aborted: the screen was blanked and a “Too Soon” on-screen error signal appeared. No visual feedback about movements was available when trials were aborted. Thus, all movements recorded and analysed were made according to the following “hard cut-off” times: within 1022.4, 272.4, 222.4, 172.4, 122.4 ms after target presentation.

**Figure 1.**
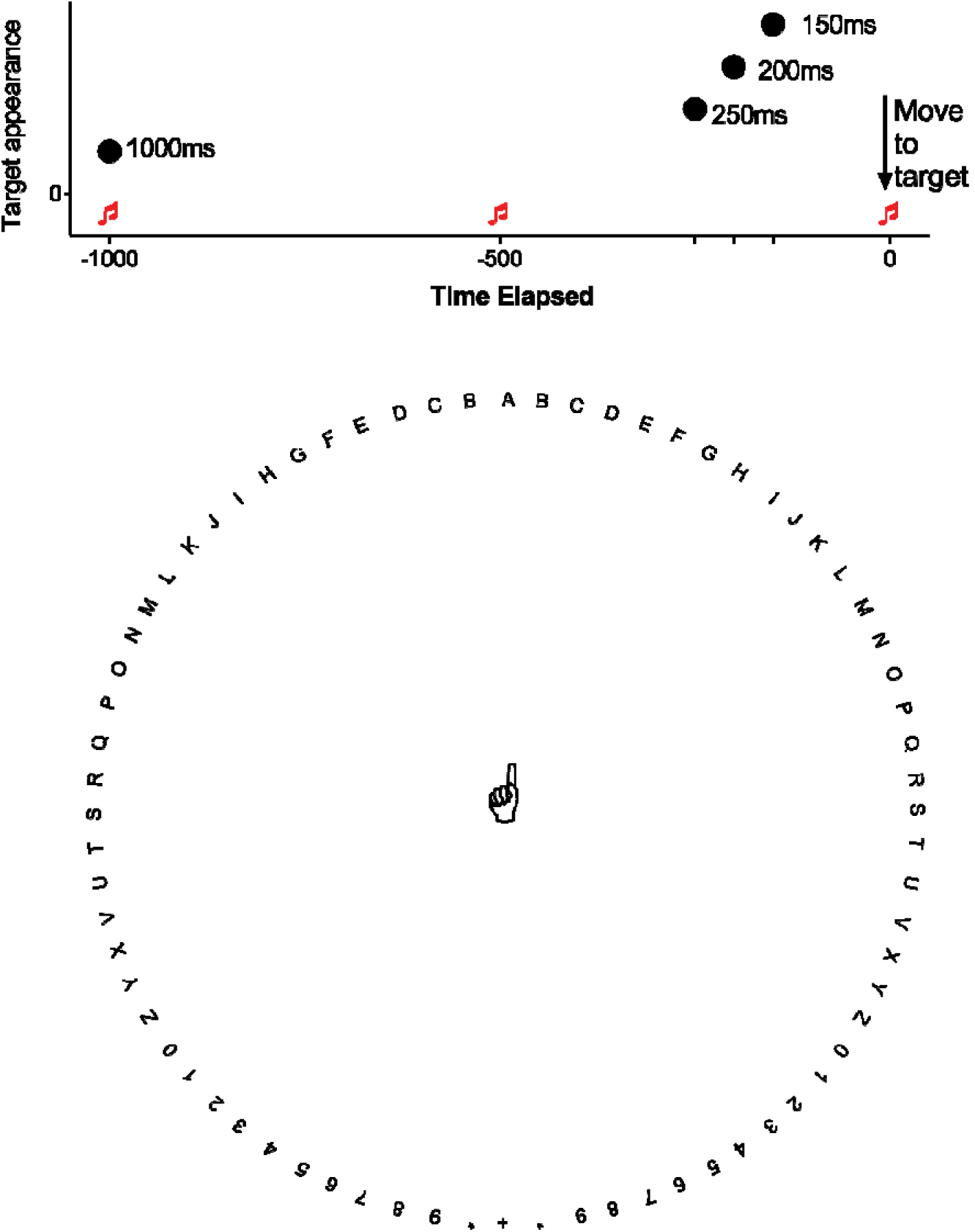
Top panel: A schematic representing the timed-response paradigm. Three tones spaced 500 ms apart were presented, and participants were instructed to time the onset of their movements with the onset of the third tone. Targets appeared at different latencies prior to the third tone (Experiment 1a: 1000ms, 250ms, 200ms, 150ms, or 100ms; Experiment 1b: 1000ms, 250ms, 200ms, 150ms; Experiment 2: Long preparation time condition: 1000ms, short preparation time condition: 250ms). Note that these latencies were minus a display latency of 27.6 ± 1.8 ms. Bottom panel: Experiment 2 landmark layout for the LongReport conditions.

#### Experiment 1

The aim was to test re-aiming performance under progressively shorter preparation times, to determine whether restricting movement preparation can prevent strategic re-aiming. This paradigm of asking participants to re-aim by a specified angle relative to a visual target is similar to that used by Georgopoulos and Massey (1987a). In each trial, participants encountered one of eight targets which either spanned a small range of 35° (−17.5°, −12.5°…17.5°) in Experiment 1A, or a distribution of 360° (0°, 45° … 360°) in Experiment 1B. Targets were presented in random order. In all trials, thirty-six “landmarks” were presented on-screen as white circles spaced 10° apart throughout the 360° range, 10 cm from the start circle. In the re-aiming condition, half of the participants were instructed to re-aim to the third landmark located clockwise from the target, and half were instructed to re-aim to the third landmark counter-clockwise to the target (i.e., 30° either side of the target). All participants completed the aiming condition before the re-aiming condition in blocks of 48 trials for each preparation time condition. The preparation times were progressively shortened, such that the trial schedule was: 1000ms aiming, 1000ms re-aiming, 250ms aiming, 250ms re-aiming, 200ms aiming, 200ms re-aiming, 150ms aiming, 150ms re-aiming, 100ms aiming, 100ms re-aiming. The 100ms condition was not included in Experiment 1B because most participants could not initiate target-directed movements prior to the deadline.

#### Experiment 2

To examine whether shortening preparation time can provide a sufficient assay of implicit learning, we compared adaptation behaviour with short preparation time to an estimate of implicit learning obtained by subtracting self-reported aiming direction from the actual direction of hand movement (Bond and Taylor 2015; Brudner et al. 2016; McDougle et al. 2015; Taylor et al. 2014). Participants were assigned either to a 250ms preparation time condition (Short), or one of two 1000ms preparation time conditions. In the LongReport condition, they had to verbally report aiming directions by stating which of 72 landmarks spaced 5° apart most closely corresponded to the direction that they were aiming towards (Bond and Taylor 2015; Taylor et al. 2014). Previous studies exclusively used numerical landmarks (Bond and Taylor 2015; Brudner et al. 2016; Morehead et al. 2015; Taylor et al. 2014), which allowed the use of mental addition or subtraction strategies in some participants (Bond and Taylor 2015). We thus avoided using only number landmarks. Landmarks consisted of the letters A to Z, the numbers 1-9, and the symbol “*” (reported as “star”). For ease of reporting, multiple-syllable characters (i.e., W) were not used. Landmarks rotated with the target, such that the same landmarks would always appear in the same location relative to the target, because rotating landmarks are more sensitive to explicit processes than fixed-location landmarks (Bond and Taylor 2015). Because of this, only a subset of the possible landmark values (A, B…G, *, 1, 2, …9) were actually used by participants when reporting their aiming directions. Participants were allowed to report their aiming direction at any time between target appearance and movement completion. Verbal reports of aiming directions were recorded online by the experimenter. To estimate implicit learning, these self-reported aiming directions were subtracted from actual movement directions. A third control group (LongNoReport) had a 1000ms preparation time, but did not have to report aiming directions. We did not apply the reporting manipulation to the Short condition, as piloting showed that it was extremely difficult to report the aiming direction when the target appeared 250 ms prior to the imperative signal to move.

Prior to the start of the experiment, participants were given no information about the nature of the rotation; they were only told that a disturbance of the cursor would be present in some trials, which may increase task difficulty. Participants in all conditions first completed a **pre-rotation** block of 6 cycles (48 trials) with veridical feedback of their movement trajectories to familiarize them with the task. LongReport participants began to verbally report their aiming direction in last 24 trials in the pre-rotation block to familiarize them with the reporting procedure. The pre-rotation block was followed by a **rotation** block (60 cycles, i.e., 480 trials) with either a 30° clockwise or counterclockwise rotation of visual feedback relative to the centre of the start circle. Halfway through this block, participants were given a 30 second break. The rotation block was followed by a **no-feedback** block of 6 cycles (i.e., 48 trials), where visual feedback of the cursor position was hidden immediately after the cursor left the start circle. Crucially, before commencing this block, participants were explicitly instructed that there was no longer any disturbance of visual feedback, and that they should aim straight towards the target (Heuer and Hegele 2008; Taylor et al. 2014). The no-feedback block therefore provides an alternative assay of implicit remapping. Finally, participants completed a **washout** block of 6 cycles (48 trials) where unrotated visual feedback was available to enable participants to return movements back to the unadapted state. Landmarks were removed from the no-feedback block and the washout block, and participants were no longer required to report aiming direction in these blocks. The same preparation time constraints were maintained throughout the entire experiment for each group.

### Data analysis

Movement onset time was taken as the time at which hand speed first exceeded 2 cm/s. Movement direction was quantified 100ms after movement onset, prior to the potential influence of online corrections. For Experiment 2, data from the counterclockwise rotation group were sign-transformed to allow us to collapse the dataset with data from the clockwise rotation group. Negatively signed angles indicate that the deviation in hand direction relative to the target was opposite to the direction of the rotation (i.e., to reduce visual error).

#### Experiment 1

To determine which of the preparation times was sufficiently short to suppress strategic re-aiming, we first quantified movement directions relative to the target as mean vectors and variability of movement directions as mean vector lengths, denoted as ***r*** for all preparation times tested using circular statistics. In the aiming condition, mean vectors values close to zero suggest that movement directions were close to the target. In the re-aiming condition, values close to 30° indicate that movement directions were close to the instructed re-aiming direction. Longer mean vectors indicate less variable movement directions, with a value of 1 indicating all directions aligned, and a value of 0 indicating an absence of directional tuning (i.e. a uniform distribution throughout all possible directions). We then compared movement directions and variability for the aiming conditions to the re-aiming conditions. When directional data is normally distributed, one can use the Hotelling’s Paired Test, which is the equivalent of the paired t-test for circular statistics (Zar 2010). However, as aiming directions were not normally distributed, we used a non-parametric alternative (Moore’s paired sample second order tests) to determine whether mean vectors differed reliably between aiming and re-aiming conditions (Zar 2010). Similarly, mean vector lengths typically show skewed distributions close to 1, and thus Wilcoxon-Rank analyses were used to compare variability between the aiming and re-aiming conditions. Circular statistics analyses were conducted with the software Oriana. For Experiment 1a (narrow target distribution), we also tested whether participants re-aimed by moving towards the middle of a (hypothetical) re-aiming target distribution by measuring the errors made to each target, for the two shortest preparation time conditions (100 ms & 150 ms). If re-aiming errors were smallest at the central 0° target and largest at the surrounding targets, then this would suggest that participants adopted a strategy to re-aim to the middle of the hypothetical re-aiming target distribution by initiating movements prior to full integration of target direction information.

#### Experiment 2

Prior to statistical analyses, movements further than 90° clockwise or counterclockwise away from the target (i.e., outside of a 180° range) were deemed as outliers, and were discarded from the analysis. This procedure excluded a small proportion of trials (Short: 4.00%, LongReport: 0.58%, LongNoReport: 0.39%). We evaluated whether the direction of hand movement relative to the target, under reduced movement preparation time conditions, was similar to the estimate of implicit learning obtained by subtracting self-reported aiming directions from actual movement directions (Taylor et al. 2014). To this end, we recoded verbal reports of landmarks into angular aiming directions, and then estimated implicit learning by subtracting reported aiming directions from actual movement directions. Trials were averaged in cycles of eight (one trial for each target angle) for statistical analysis. To compare adaptation behaviour between conditions, ANOVAs with the within-subjects factor Cycle and two between-subjects factors of Condition and Rotation Direction (clockwise, counterclockwise) were run on relevant cycles. For the early adaptation phase, the relevant cycles were cycles 1-30 of the adaptation block. For the late adaptation phase, the relevant cycles were cycles 31-60 of the adaptation block. For the no-feedback block, the relevant cycles were all 6 cycles of the no-feedback block. For the washout block, the relevant cycles were all 6 cycles of the washout block. For all ANOVAs, when Mauchly’s test of sphericity was statistically significant, the Greenhouse-Geisser correction was used to adjust degrees of freedom.

## Results

### Experiment 1: Re-aiming away from a target at very short preparation times

Movement directions for all trials pooled across all subjects are shown in Figure 2 for Experiment 1A (small target range) and for Experiment 1B (large target range). With the small target range, movement directions were close to the target directions when aiming, and approximated the required 30° offset when re-aiming, even with the shortest preparation time condition of 100ms (i.e., hard initiation cut-off of 122.4ms). Rao’s tests run for each participant’s dataset within each preparation time condition indicated that movement directions were directionally tuned for all conditions, even for the shortest 100ms preparation time condition (all p<.0001).With the large target range, re-aiming movements were directed progressively closer to the original target (i.e., further from the instructed 30° offset) as preparation times were shortened. Rao’s tests run for each participant’s dataset within each preparation time condition indicated that movement directions were not directionally tuned for 5 of the 13 participants who completed the 150ms aiming condition and 10 of the 13 participants who completed the150ms re-aiming conditions.

**Figure 2.**
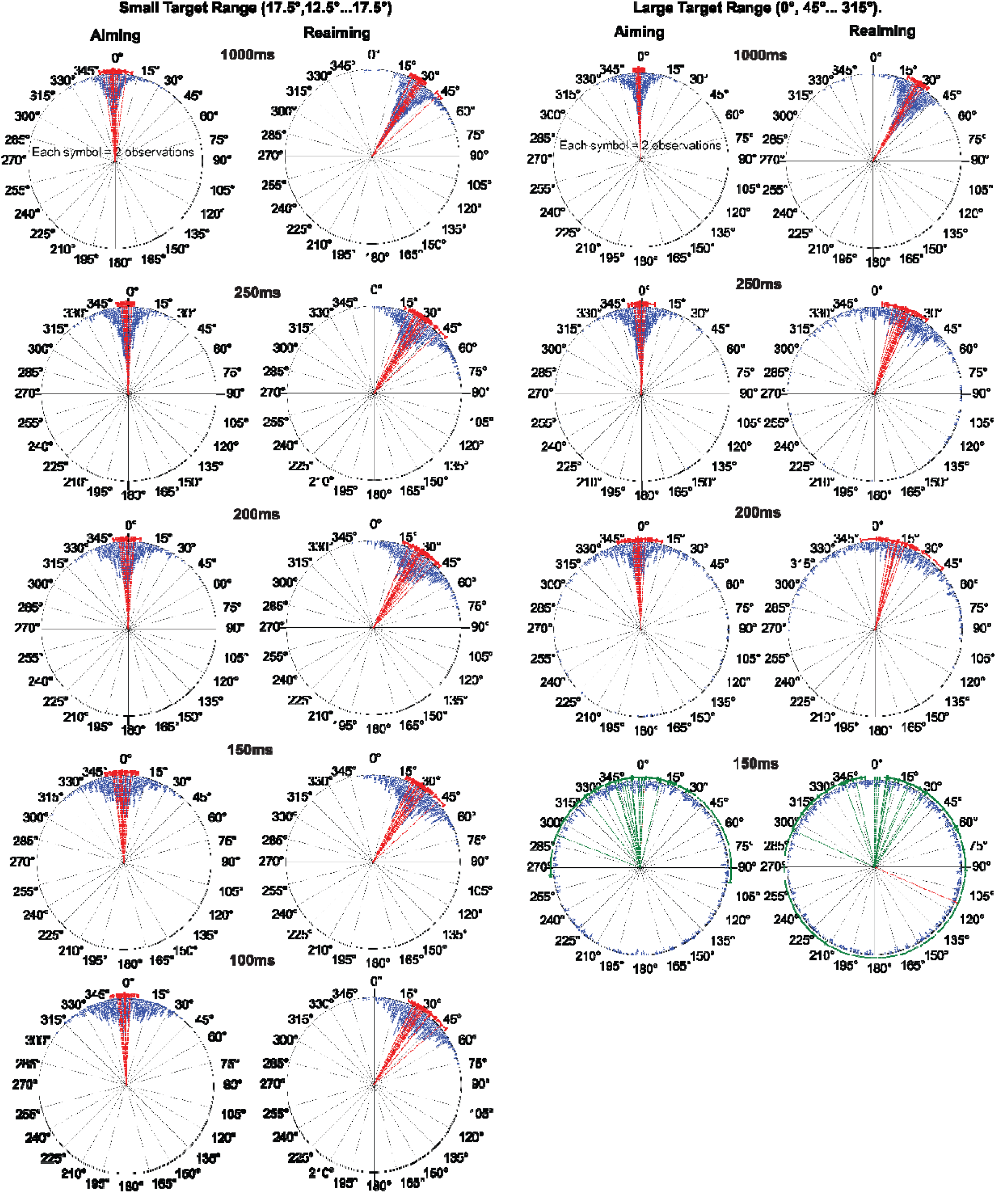
Movement directions for the narrow target range (−17.5° to 17.5°) and large target range (0° to 360°) plotted relative to target direction at 0°, in the aiming and re-aiming conditions. Data from participants in the counterclockwise re-aiming condition were normalized to the clockwise direction and collapsed with data from participants in the clockwise re-aiming condition. Symbols represent movement directions in individual trials for all participants across the preparation time conditions (1000ms, 250ms, 200ms, 150ms to 100 ms). Note that the hard cut-off times for movement initiation in these conditions were: 1022.4, 272.4, 222.4, 172.4, 122.4 ms after target appearance. Red vectors represent individual mean vectors for each participant, and error bars represent the mean and 95% confidence intervals of mean movement direction for each participant. Green vectors represent individual mean vectors that were not significantly directionally tuned according to a Rayleigh’s test.

Table 1 summarizes statistical comparisons between aiming and re-aiming across preparation times for both the narrow target distribution (Experiment 1A) and the full 360° target distribution (Experiment 1b). For both target distributions, movement directions were more variable (shorter vector lengths) when re-aiming away from the target than when aiming to the target across all preparation times. When errors were averaged across all targets in the narrow range (Experiment 1a), it appears that participants could re-aim away from the target in all preparation times tested (even when movements were initiated within 122.4 ms of target preparation). That is, mean vector angles were on average approximately 30° away from the target across all preparation times tested. We were surprised at this apparent success in re-aiming, since Haith et al (2016) showed that directionally tuned movements to a unique target require approximately 130 ms, and the process of re-aiming (and preparing movements to a direction offset by 30° to the target) should require at least some additional processing (Haith et al. 2016). We therefore examined errors for each target in the distribution individually, to search for evidence that participants might have been able to achieve task success by aiming toward the middle of the re-aiming target distribution (i.e. 30° away from the central visual target distribution). In this case, movement could be initiated prior to integration of target direction information, but average errors collapsed across targets would be close to zero.

**Table 1:**
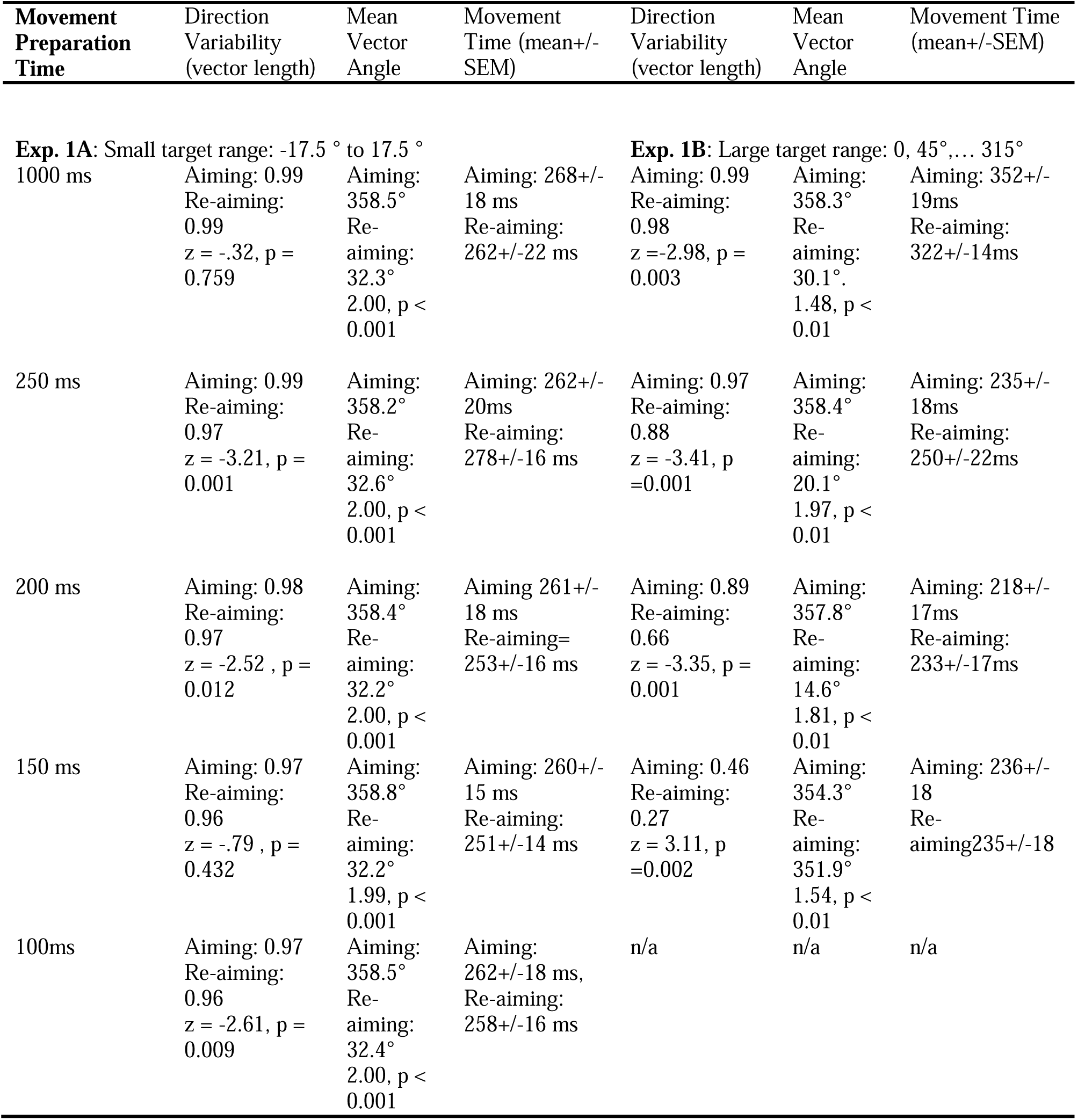
Statistical analyses comparing aiming and re-aiming accuracy (assessed via mean movement direction) and variability (assessed via vector length—longer vectors indicate less variability), as the amount of time available to prepare movements was progressively shortened.

Figure 3 shows clear evidence that subjects adopted such a strategy for the shortest preparation time condition, under both aiming and re-aiming conditions. Errors were similar for all targets in the 150ms preparation time condition, indicating that there were no large inherent biases in reaching performance. There were no statistically significant differences in error size across targets (F(7,91) = 1.10, p = 0.39, partial η-squared = 0.08) or conditions (F(1,13) = 1.1, p = 0.3, partial η-squared = 0.08), nor an interaction between target and condition (F(7,91) = 1.2, p = 0.3, partial η-squared = 0.09). By contrast, with 100ms preparation time (122.4 ms hard cut-off), errors were systematically larger in absolute terms as the angle from the centre of the distribution increased for the aiming condition (main effect of target F(7,91) = 199, p < 0.001, partial η-squared = 0.94). The signs of errors indicate that participants made reaching movements that were biased towards the central target. The pattern of errors for aiming and re-aiming conditions were similar for the aiming and re-aiming conditions, with no statistically significant main effect of condition (F(1,13) = 0.6, p = 0.45, partial η-squared = 0.04) or interaction between condition and target (F(7,91) = 1.6, p = 0.15, partial η-squared = 0.11). Note that errors from the required (re-aiming) target are plotted and analysed, rather than errors relative to the presented target. Critically, the similarity in error directions and magnitudes for the aiming and re-aiming conditions, across all preparation time conditions, suggests that if participants had sufficient time to aim towards the target, then they also had time to re-aim to one side of the target by a specified angle. Although this process of re-aiming must require some additional processing, our data suggest that such processing is extremely rapid, to the point that we were not able to detect a time-cost for re-aiming under the conditions of our experiment. The data also suggest that people are able to apply a re-aiming strategy to an anticipated target location when there is insufficient time to adequately process visual information related to the actual target. This indicates that the approach of restricting strategic re-aiming through preparation time constraints might be especially problematic for single or dual target paradigms.

**Figure 3.**
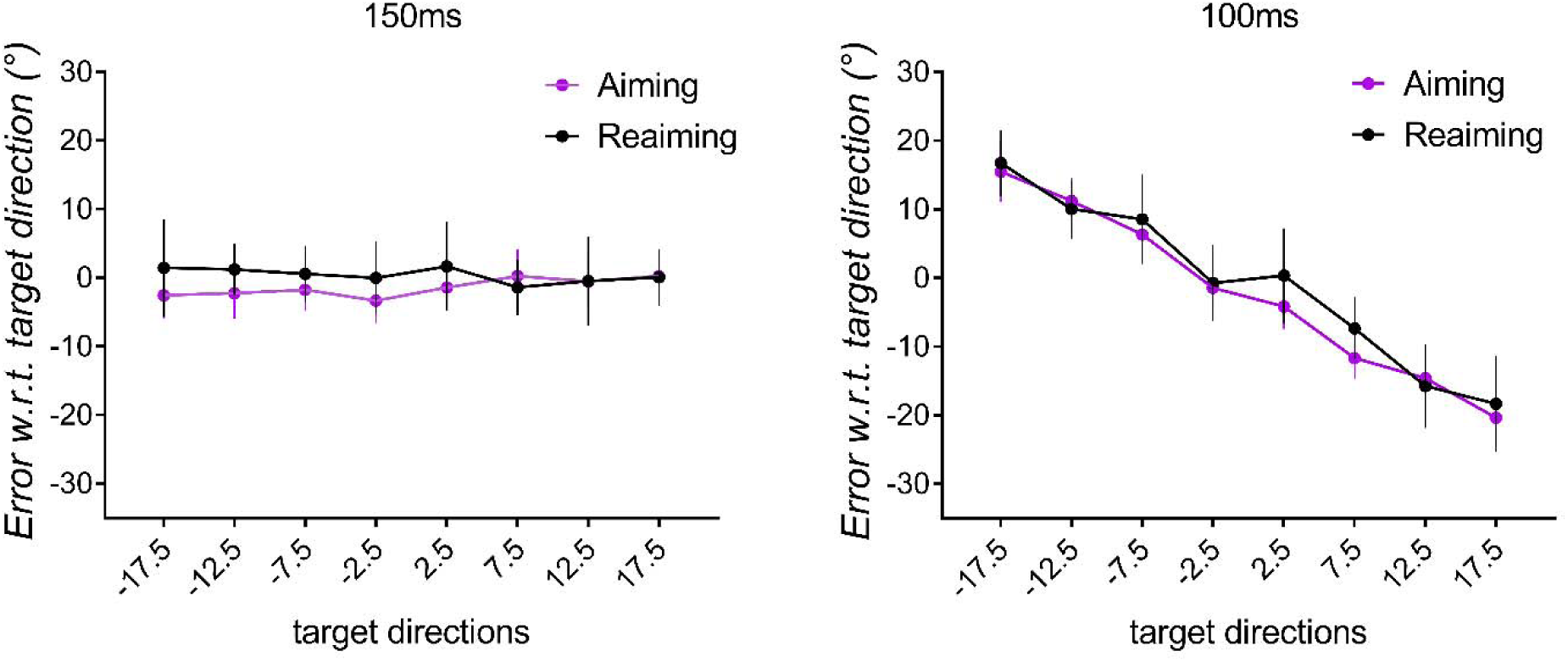
Movement errors for each target direction from −17.5° to 17.5° with respect to (w.r.t) the required reaching direction (i.e., presented target or re-aiming target depending on condition). Data from participants in the counterclockwise re-aiming condition were normalized to the clockwise direction and collapsed with data from participants in the clockwise re-aiming condition. Separate plots are shown for the 150ms to 100 ms preparation time conditions. Note that the hard cut-off times for movement initiation in these conditions were 172.4 and 122.4 ms after target appearance. Values are group mean errors and error bars represent 95% confidence intervals.

In Experiment 1b, which involved the broad target distribution, participants were less accurate at re-aiming away from the target (20.1°) with 250ms preparation, although re-aiming away from the target was still possible with 200 ms (14.6°) and 150ms (7.7°) preparation. This confirms that voluntary re-aiming is not absolutely prevented by shortening movement preparation time, irrespective of whether potential targets lie within a narrow or large angular range. Self-reports from our participants indicated, however, that re-aiming was extremely effortful at short preparation times, especially when targets were distributed around the circle. Moreover, the accuracy cost of re-aiming was dramatically greater when targets were distributed around the circle. Given this, in Experiment 2, we considered whether participants would choose to re-aim under time pressure in order to improve performance in a visuomotor rotation task. For this experiment, targets were radially arranged throughout the circle (0°, 45°…315°) and movement preparation time was restricted to 250ms. We decided to use 250ms as an arbitrary trade-off between a sufficient time to allow accurate aiming to the presented target, and sufficient time-pressure to make re-aiming effortful.

### Experiment 2: Suppressing strategic re-aiming with short preparation time constraints reduces the rate and extent of error compensation

Figure 4 shows the group mean, cycle-averaged, movement directions across different phases of the experiment. To evaluate whether the discrepancy between the measures of implicit learning (i.e., implicit learning estimated from subtracting aiming directions from movement directions and implicit learning estimated from the no-feedback trials) is related to the process of reporting explicit aiming angles or the preparation time constraints, we compared this data to 10 additional task-naïve participants (5 counterclockwise, 5 clockwise) who completed the visuomotor rotation task with the same 1000ms preparation time constraints via the same timed-response paradigm, but who did not report aiming directions and had no visual landmarks throughout the task (LongNoReport). In the baseline block (i.e., before encountering the perturbation) a counterclockwise bias was evident in the Long preparation time group, as Cycle (Cycle 1…6) x Condition (Long, Short, LongNoReport) x Rotation Direction (clockwise, counterclockwise) ANOVA revealed a significant main effect of Condition, F(2,30) = 4.267, p = 0.023, partial η-squared = 0.221. To estimate the bias, we averaged mean movement directions from baseline cycles 2-6 (baseline cycle 1 was not included as participants were still familiarising themselves with the vBOT at this stage). To eliminate the influence of this bias on the subsequent test phases, we subtracted the bias from mean movement directions from each subsequent cycle (i.e., the first cycle of the adaptation block to the last washout cycle). The adaptation phase was arbitrarily separated into Early (Cycle 1-30) and Late blocks (Cycle 31-60). ANOVAs were run on each block for all three conditions (LongReport, Short, LongNoReport), according to a mixed within-between effects model (Cycle x Rotation Direction x Condition [LongReport, Short, LongNoReport]). In the Early phase, there was a significant main effect of Condition, F(2,30) = 6.25, p = 0.005, partial η-squared = 0.294, as well as a significant Cycles x Condition interaction, F(24.6,370.2) = 1.59, p = 0.037, partial η-squared = 0.09, as less error compensation was evident with Short (−17.3+/−1.3°) compared to LongReport, (−22.4+/−1.3°, p=.033) and compared to LongNoReport (−24.4+/−1.8°, p=.009). Error compensation in this early phase did not differ reliably between LongReport and LongNoReport (p=.75). Similarly, for the Late phase, there was a significant main effect of Condition, F(2,30) = 4.77, p = 0.016, partial η- squared = 0.241; as less error compensation was evident with short preparation time (−23.1+/−1.1°) compared to LongNoReport (−28.7+/−1°, p = .007) and compared to LongReport (−26.7+/− 1.1°, p =.036). Error compensation was also more complete for clockwise than counterclockwise rotations, as evident in significant main effect of Rotation across all phases: Early: F(1,30) = 21.643, p < 0.001, partial η-squared = 0.419, Late: F(1,30) = 10.96, p = 0.002, partial η-squared = 0.268]. There were no other significant interactions.

**Figure 4.**
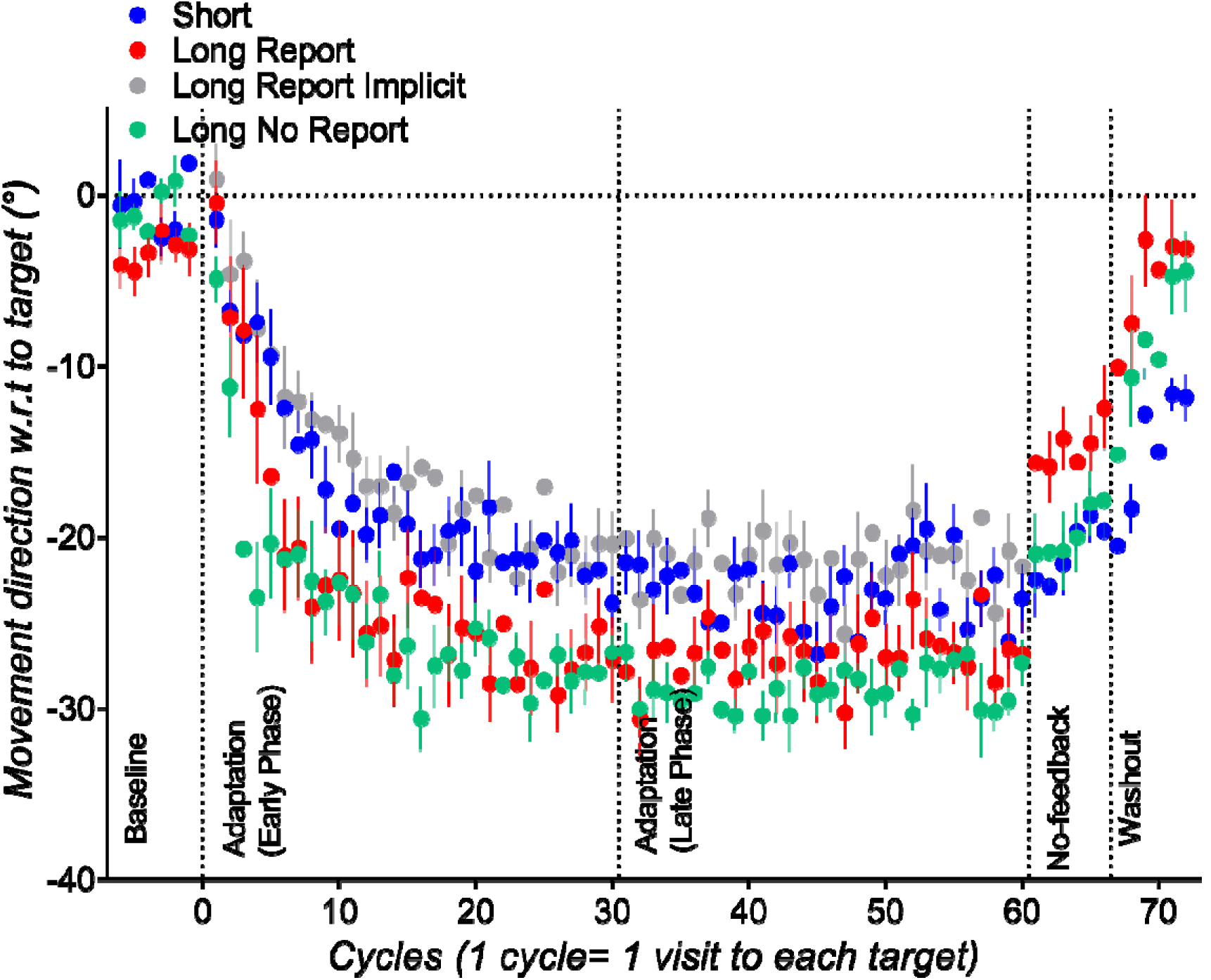
Experiment 2 mean movement direction in every cycle, averaged across each condition. Data from participants who encountered counterclockwise rotations were sign-transformed to allow collapsing with data from participants who encountered clockwise rotations. Error bars are standard errors of the mean. Negative values indicate movements that were opposite from the direction of rotation, positive values indicate movements that were in the same direction as the rotation. Note that Long Report Implicit is not an additional experimental condition, but is derived from subtracting self-reported aiming directions from movement directions in the Long Report condition.

### Preparation time constraint as an assay of implicit learning

The implicit component of error compensation observed for the Long preparation group was estimated by subtracting the participants’ reported aiming direction from their actual movement direction, similar to previous work (Bond and Taylor 2015; Brudner et al. 2016; McDougle et al. 2015; Taylor et al. 2014). This measure of error is hereafter termed “LongImplicit”, and was compared to angular errors observed between the target and movement for the short preparation time condition. There were no significant differences between LongImplicit and Short, as shown by Condition (LongImplicit, Short) x Cycle (Cycle 1…30) x Rotation Direction (CW, CCW) ANOVAs run for the early adaptation phase [main effect of Condition, F(1,24) = 1.33, p = 0.26, partial η-squared = 0.05, Cycle x Condition F(12.6,303.9) = 1.05, p = 0.4, partial η-squared = 0.04 interaction], as well as the late adaptation phase [Condition, F(1,24) = 1.44, p = 0.2, partial η-squared = 0.06, Cycles x Condition, F(11.9,287.1) = 1.4, p = 0.16, partial η-squared = 0.05]. The main effect of rotation direction was statistically significant for the early adaptation phase, F(1,24) = 26.29, p < 0.001, partial η-squared = 0.52 as well as for the late adaptation phase: F(1,24) = 11.473, p = 0.002, partial η-squared = 0.32. There were no significant interactions. Thus, the extent and rate of implicit learning did not differ reliably between estimates based on subtracting self-reported aiming directions and restriction of preparation time.

### Difference in estimate of implicit learning from subtracting aiming directions and estimate of implicit learning from no-feedback trials

An alternative measure of implicit remapping is provided by the no-feedback trials that participants performed after the final adaptation phase block. Here, participants received no visual feedback about their movements, and were explicitly instructed that the perturbation was removed and that they should aim straight to the target (Taylor et al. 2014), (similar to Heuer and Hegele 2015). For the LongReport group (Figure 3) the measure of implicit learning obtained from this no-feedback block appears substantially lower (i.e., movements were less adapted) than the measure of implicit learning obtained by subtraction of reported aiming direction in the last adaptation cycle. In contrast, for the Short group, errors in the last adaptation cycle were similar to those in the first no-feedback cycle. To compare implicit learning (estimated by subtracting aiming direction or by shortening preparation times) to implicit learning estimated by no-feedback trials, we compared the last adaptation cycle (after subtracting aiming directions for the LongReport group) to the first no-feedback cycle for the LongReport group and the Short group, via a Condition (LongReport, Short) x Rotation Direction (CW, CCW) x Phase (last adaptation cycle, first no-feedback phase cycle) ANOVA. There was a significant Phase x Condition interaction, F(1, 24) = 4.36, p = .047, partial eta-squared= .15. Follow-up Rotation Direction x Phase (last adaptation cycle, first no-feedback phase cycle) ANOVAs were run separately for the LongReport and the Short condition. For LongReport, implicit learning estimated by subtracting aiming direction in the last adaptation cycle (21.7+/−1.8°) was more than implicit learning estimated in the first no-feedback cycle (15.8+/−1.6°), as shown by a significant main effect of phase F(1,12) = 6.94, p = 0.022, partial η-squared = 0.37. In contrast, for the short preparation time, the last adaptation cycle (−23.5+/−1.8°) did not differ reliably from the first no-feedback cycle (−22.5+/−1.9°): the main effect of Phase was not significant (F(1,12) = 0.33, p = 0.57, partial η-squared = 0.02), and did not interact significantly with any other factor. Thus, for the LongReport group, there was a discrepancy between the estimates of implicit learning provided by the reporting method, obtained in the presence of the rotation, and the no-feedback condition, obtained after the final movement performed under the visuomotor rotation. There was no discrepancy between implicit learning estimates for the short preparation time group, even though the final estimate of implicit learning at the end of adaptation was similar to that obtained after subtracting aiming directions for LongReport group, and despite the fact that both groups had explicit knowledge that the rotation was removed.

This discrepancy between the estimates of implicit learning from reporting, in the last adaptation cycle, and from no-feedback trials in which participants were instructed that the rotation was absent, was also evident in previous work using the reporting procedure (c.f. Fig 2C, Fig 5C Bond and Taylor 2015). Taylor et al. (2014) attributed the effect to trial-by-trial decay of adaptation within the first no-feedback cycle, because there was no statistically significant difference between the last adaptation trial and the first no-feedback trial (Taylor et al. 2014). Our LongReport group similarly showed no reliable difference in estimated implicit learning from the last adaptation trial to the first no-feedback trial (Trial x Rotation Direction ANOVA on the LongReport group showed a non-significant main effect of Trial F(1,12)=.30, p =.59, partial eta-squared =.03). However, we hesitate to make inferences from this non-significant effect, because comparing trial-by-trial data in multi-target designs can be problematic: target directions were likely to differ between the last adaptation trial and the first no-feedback trial between-subjects, and directional accuracy differs between targets (Gordon et al. 1994). Moreover, movements were also less adapted on average over all six no-feedback cycles for the LongReport than the Short group, as shown by a significant main effect of Condition, F(1,24) = 6.91, p = 0.01, partial η-squared = 0.22 in a Condition x Rotation Direction x Cycle ANOVA. This suggests that the extent or persistence of implicit learning was less for the long preparation with reporting condition than the short preparation condition.

To evaluate whether the discrepancy between measures of implicit learning is related to the reporting procedure (i.e., the process of reporting explicit aiming angles and/or the presence of visual landmarks), we compared error compensation data from the Long Report group to the LongNoReport group. Error compensation during exposure to the rotation did not differ reliably between this LongNoReport group and the LongReport group, as Cycle x Reporting (LongNoReport, LongReport) x Rotation Direction (CW, CCW) ANOVAs run separately for the early adaptation phase (Cycles 1…31) and the late adaptation phase (Cycles 31…60) showed a non-significant main effect of reporting for the early adaptation phase [F(1,18) = 0.67, p = 0.424, partial η-squared = 0.036], and no significant interactions, as well as for the late adaptation phase, F(1,18) = 0.843, p = 0.371, partial η-squared = 0.045, no significant interactions]. However, the estimate of implicit learning obtained from no-feedback trials was greater for the LongNoReport group than the LongReport condition: Cycle (Cycle 1-6) x Reporting (LongNoReport, LongReport) x Rotation Direction (CW, CCW) ANOVA on the no-feedback block showed a significant main effect of reporting, F(1,18) = 7.32, p = 0.015, partial η-squared = 0.289. There were no other significant interactions. The main effect of Rotation Direction was significant F(1,18) = 16.64, p = 0.001, partial η-squared = 0.48—similar to the adaptation phase, movements were more adapted with the clockwise direction (−21.0+/−1.0°) than the counterclockwise direction (13.4+/−1.0°). Performance in the no-feedback trials did not differ significantly between the LongNoReport and the Short group—a Cycle (Cycle 1-6) x Condition (LongNoReport, Short) x Rotation Direction ANOVA showed a non-significant main effect of condition [F(1,18) = 0.449, p = 0.511, partial η-squared = 0.024], and no significant interactions with condition, all p>0.5. The main effect of rotation direction was significant F(1,18) = 15.98, p = 0.001, partial η-squared = 0.47.

## Discussion

This study aimed to evaluate a previously established method of assaying implicit learning by restricting the time available to prepare movement (Fernandez-Ruiz et al. 2011; Haith et al. 2015). Experiment 1 showed that restricting time available to prepare movements does not prevent people from applying a deliberate strategy to re-aim to one side of a target, particularly when targets are distributed within a narrow angular range. However, Experiment 2 showed that restricting movement preparation time effectively reduces strategic re-aiming during adaptation to visuomotor rotation when targets are distributed throughout 360°, as shown by slower and less complete error compensation compared to when movement preparation times were not shortened. Moreover, the errors made by participants when preparation time was shortened were indistinguishable from an assay of implicit learning obtained by subtracting self-reported aiming directions from movement directions (Bond and Taylor 2015; Brudner et al. 2016; McDougle et al. 2015; Taylor et al. 2014). Surprisingly, despite this similarity in estimates of implicit learning obtained for the two methods during exposure to the visuomotor rotation, participants who reported aiming directions showed less implicit remapping in the post-perturbation no-feedback trials than those who did not report aiming directions. This suggests that the process of reporting aiming direction reduces the extent or persistence of implicit learning.

### Suppressing the expression of explicit learning by restricting preparation time

Despite a long history of studies on implicit and explicit processes in sensorimotor adaptation (Jakobson and Goodale 1989; Uhlarik 1973), our understanding of how these processes interact to determine behaviour remains incomplete. Here, we further evaluated the method of assaying implicit learning by restricting movement preparation time (Fernandez-Ruiz et al. 2011; Haith et al. 2015). We showed that when there is intention to re-aim (i.e., when participants were explicitly instructed to re-aim) and potential targets were distributed within a small (35°) range, accurate re-aiming is possible irrespective of the time between target presentation and movement initiation. The accuracy cost of re-aiming in such conditions was modest. Moreover, for the shortest preparation time condition (movement initiation constrained to occur within 123 ms of target presentation), it appears that participants initiated movement prior to complete integration of visual information about the actual target, and were able to achieve task success by aiming or re-aiming to the centre of the (required) target distribution. When target direction (and thus re-aiming direction) was less predictable (targets distributed throughout 0-360°), however, re-aiming accuracy declined with progressively shorter preparation times. Participants were still able to partially re-aim away from the target whenever they had sufficient time to produce directionally tuned movements, but at the expense of dramatically increased movement variability. Hence, compressing preparation time does not introduce an absolute limit upon the capacity for re-aiming, particularly for narrow target distributions.

However, during sensorimotor adaptation to a perturbation, restricting preparation time appeared to suppress re-aiming when targets were distributed about 360°, such that error compensation was indistinguishable from the assay of implicit learning obtained from subtracting reported aiming direction from actual movement direction. This suggests that people choose not to apply re-aiming strategies to correct for visuomotor perturbations under time pressure, presumably to avoid the increases in effort and variability associated with re-aiming under such conditions.

This interpretation prompts a formal definition of the distinction between implicit and explicit processes. Here, consistent with others (Huberdeau et al. 2015), we define explicit processes as those which can be deliberately engaged and disengaged. By contrast, implicit processes are automatic and difficult to deliberately disengage. We do not distinguish between explicit processes from implicit processes based on awareness of the perturbation or a re-aiming strategy, as classically defined (Reber 1967). Indeed, many of our participants in the short preparation time condition were able to accurately describe the nature of the rotation and could articulate a compensatory strategy, but found it simply too difficult to implement the strategy when preparation times were restrained.

Our findings that asymptotic error levels were greater for short than long preparation time conditions differ from those of Haith et al. (2015). In their task, which involved two potential targets, participants were eventually able to reduce errors to a similar degree for the short and long preparation time targets. This discrepancy in findings probably relates to the predictability of the target locations. Targets only appeared in two locations in Haith et al. (2015), with preparation time of ~300 ms. However, our Experiment 1A shows that explicit re-aiming is possible even at 123 ms when the target direction was predictable within a small 35° range. Hence, although the target-switch protocol in Haith et al. (2015) appears to have restricted explicit processes initially, the method may not have been sufficient to suppress re-aiming by the end of the adaptation block.

### Discrepancy between different estimates of implicit learning

In Experiment 2, the extent of implicit learning inferred from aiming reports in the long preparation time condition was similar to the extent of error compensation observed for the short preparation time condition. However, for the long preparation condition, there was a difference between estimates of implicit learning obtained from reporting during exposure to the rotation, and estimates of implicit learning obtained from subsequent movements made without feedback. A discrepancy has been reported previously between measures of implicit learning measured via movement directions after subtracting aiming directions, and via movement directions in subsequent no-feedback trials (c.f. Fig 2C, Fig 5C Bond and Taylor 2015). However, we found that there was no such decay between errors in the last perturbation trials and first no-feedback trials for the short preparation time condition. Furthermore, the overall amount of implicit remapping (indicated by adapted movements in the no-feedback block despite explicit knowledge that the rotation had been removed),was less in the reporting group than in either of two groups that did not report aiming directions (i.e., the LongNoReport group and the Short group), irrespective of movement preparation time. We note that this difference might result from either the act of reporting aiming directions, and/or the presence of visual landmarks, however, as the original reporting procedure often requires the use of landmarks, we did not attempt to dissociate between the two possibilities.

We propose two possibilities to account for these observations. One possibility is that implicit learning is more labile (i.e., more sensitive to decay due to a change in task context or the passage of time) when it is acquired in a context in which people report their re-aiming strategies to compensate for errors. The proposal that explicit processes reduce the persistence of implicit remapping is consistent with findings in prism adaptation, where explicit knowledge of the nature of the perturbation reduces the extent of implicit remapping measured in post-perturbation no-feedback trials (Jakobson and Goodale 1989; Uhlarik 1973). One caveat to this interpretation is that, although all three groups experienced the same change in context (i.e., from having feedback of cursor position with visuomotor rotation to having no cursor feedback and explicit knowledge that the rotation had been removed), the LongReport group experienced an additional context change (i.e., from having to report aiming directions to no longer having to report aiming directions). Thus, we cannot rule out the possibility that the extent of context change, rather than sensitivity to change, was the key factor underlying a reduced estimate of implicit learning in the LongReport condition.

An alternative possibility that could explain our data is that people may have systematically under-reported their aiming angle (i.e., people re-aimed to a greater extent than they reported). This would result in an underestimation of explicit learning and an overestimation of implicit learning in the error compensation phase. In this case, the no-feedback trials would provide a more accurate measure of implicit learning than the reporting trials, which in turn would imply that the reporting procedure enhanced explicit learning and impaired implicit learning relative to non-reporting conditions. The possibility that the reporting procedure enhanced explicit re-aiming is supported by previous findings of faster error compensation with the reporting procedure than without (Taylor et al. 2014). Such a situation would suggest a competitive push-pull relationship between implicit and explicit processes in sensorimotor adaptation. A push-pull relationship between implicit and explicit processes has been shown for other motor learning tasks. For example, in sequence learning, disrupting explicit awareness of a sequence to be learned, by performing a concurrent verbal declarative task, improved post-task recall of implicitly acquired sequences (Brown and Robertson 2007). Similarly, in force-field adaptation, engaging a declarative verbal memory task resulted in poorer recall of a fragile, possibly explicit memory created by a fast process, and improved recall of a robust, possibly implicit memory created by a slow process (Keisler and Shadmehr 2010).

By contrast, implicit adaptation to visuomotor rotation has been argued to be inflexible, such that it develops in parallel with, but independently from, explicit learning (Bond and Taylor 2015). Although it is difficult to test whether self-reports of aiming direction are accurate, discrepancies between self-reported aiming directions and actual aiming directions seem possible. Georgopoulos and Massey (1987a) showed that when participants were explicitly instructed to re-aim by a specified angle, their re-aiming was in excess of the instructed angle, particularly with smaller instructed re-aiming angles of less than 35°. Thus, the question of whether implicit and explicit processes operate independently or competitively in visuomotor rotation learning warrants further attention.

### Summary

This study evaluated the method of dissociating implicit and explicit learning by manipulating the amount of time available to prepare movements. The method has previously been shown to unmask implicit visuomotor rotation learning on a trial-by-trial basis (Haith et al. 2015). We found that although shortening preparation time does not prevent people from voluntarily aiming to one side of a target, it appears sufficient to suppress strategic re-aiming during visuomotor adaptation when targets are distributed about a broad angular range. Estimating implicit learning by subtracting aiming directions from movement directions yielded a discrepancy between the estimate of implicit error compensation obtained during exposure to the perturbation, and the estimate of implicit learning obtained from post-perturbation trials without feedback. It is possible that the reporting procedure inadvertently increased explicit re-aiming and decreased implicit learning, which would suggest a push-pull relationship between explicit and implicit learning. In contrast, shortening movement preparation time did not result in a discrepancy between the estimate of implicit learning obtained from self-report during exposure to the perturbation, and the estimate of implicit learning obtained from trials performed subsequently without visual feedback.

